# Batch, Prefetch, Skip, Orient, Squeeze: Faster, Smaller *k*-mer dictionaries via the Spectral Burrows-Wheeler transform

**DOI:** 10.64898/2026.11.09.765318

**Authors:** Anastasia C. Diseth, Jarno N. Alanko, Joel Mackenzie, Simon J. Puglisi

**Affiliations:** University of Helsinki, Helsinki, Finland; The University of Queensland, Queensland, Australia

## Abstract

Looking up *k*-mers in an index is a core operation in genomics, underlying read mapping, metagenomic classification, and many other tasks. Queries are often *streaming* : the *k*-mers come from a longer sequence such as a sequencing read, and consecutive *k*-mers overlap. The fastest index for such queries is currently the hashing-based SSHash; indexes based on the spectral Burrows–Wheeler transform (SBWT) are competitive in space but slower. We show how to make SBWT indexes both faster and smaller. First, we process queries in small batches and prefetch the memory each will need next, hiding the latency of the dependent random accesses that dominate a lookup; together with a new rank structure that reads one cache line per query, this makes *k*-mer lookups about seven times faster than unbatched SBWT search. Second, we extend batching to streaming search and show that one failed lookup can rule out many *k*-mers of a read at once, which removes the need for the longest common suffix array. Third, we build an SBWT that stores only one orientation of each *k*-mer, thereby halving its size, with a modified search that exploits how rarely the orientation changes along a read. Finally, we adapt a compact subset rank structure to batched search. On the collections of the SSHash benchmark, and on two collections up to 15 times larger, our fastest index answers streaming queries for 31-mers faster than SSHash on every read set, by up to 4.9 times, using 12 bits per canonical *k*-mer against SSHash’s 7.4–11.1. We also show that a 6.1-bit index is still faster than SSHash on most read sets, and that our smallest index needs about 4 bits per canonical *k*-mer. Unlike that of SSHash, the space of our indexes barely varies across collections. Our code is available at https://github.com/anadis504/sbwt-squeeze-rs.

## 1 Introduction

Looking up *k*-mers, i.e. substrings of length *k*, in an index is one of the most fundamental operations in sequence analysis. As such, it pervades modern genomics pipelines, from read mapping [24, 11] and alignment-free sequence comparison [29] to metagenomic classification [35, 3] and genome assembly [23]. Lookups arise in two broad patterns. In some applications, the queries are individual *k*-mers with no particular relationship to one another. More often, however, the query *k*-mers are not independent: they arrive embedded in a longer sequence, such as a sequencing read, so that consecutive *k*-mers overlap in *k* − 1 characters. We refer to the former as *simple queries* and to the latter as *streaming queries*.

Research on efficient methods for simple and streaming *k*-mer lookup is vast and remains intense [13, 30, 31, 26, 32, 6]. The current leading method is SSHash [30, 32], which partitions the *k*-mers into buckets according to their minimizers, stores the corresponding super-*k*-mers contiguously in a compact encoding, and uses a minimal perfect hash function over the minimizers to identify the single small bucket in which a query *k*-mer must be sought. Streaming queries benefit further because consecutive *k*-mers of a read usually share a minimizer, and hence a bucket. Other hashing-based *k*-mer lookup methods, largely superseded by SSHash, include Pufferfish [7], Bifrost [19], and BLight [25]. Another family of *k*-mer lookup methods is derived from the Burrows–Wheeler transform [12]. These include the BOSS data structure [10, 9, 23, 27] and its simplifying generalization, the spectral Burrows–Wheeler transform (SBWT) [2], whose streaming queries can be accelerated with a longest common suffix array [1]. These methods also yield small and fast *k*-mer indexes, and remain competitive with SSHash in space [32], but currently lag behind it in query time. This work reshapes the current landscape by describing SBWT-based *k*-mer lookup methods that are substantially smaller and faster than previous SBWT-based indexes, and that also outperform SSHash. For example, on the HumGut reference collection [18], our fastest method answers streaming queries 1.5–2.0 times faster than SSHash while using 12 bits per canonical 31-mer, compared with 11 bits for SSHash. Another of our methods uses just 6 bits per canonical 31-mer, little more than half the space of SSHash, and still answers streaming queries about as fast or faster. Our indexes are also predictable in size: their space depends almost only on the number of *k*-mers indexed, whereas that of SSHash varies by a factor of three across our collections and values of *k* (Section 8.3). Our contributions, in more detail, are as follows.

1. We show that the latency of SBWT lookups can be hidden by processing queries in small batches and prefetching the memory each query will need next. A single lookup is a chain of dependent, essentially random memory accesses that leaves most of the CPU’s capacity for outstanding cache misses unused; following work by Groot Koerkamp on the FM-index [17], mini-batching puts this capacity to use and answers simple queries about five times faster (Section 3).
2. We describe a rank structure, *one-line rank*, in which every query reads a single 64-byte cache line, half as many as rank9, at a similar space overhead of 23.1% (Section 3.1). Furthermore, it is not specific to the SBWT and may be of independent interest.
3. We extend mini-batching to streaming queries, which are harder to batch because a failed extension sends a read into a data-dependent sequence of back-off steps. We give batched streaming searches for both kinds of streaming support the SBWT can use, each organized around how its reads diverge (Section 4).
4. We observe that when the search of a *k*-mer fails after *l* characters, none of the *k* − *l* + 1 *k*-mers of the read containing those characters can be in the index, and we exploit this in a streaming search we call `forward-skip` (Section 5). It makes the longest common suffix array unnecessary at little or no cost in speed, shrinking the index and its streaming support from 20 to 12 bits per canonical *k*-mer.
5. We show how to build an SBWT that holds only one orientation of each *k*-mer, rather than each *k*-mer together with its reverse complement, reorienting unitigs to minimize the number of dummy *k*-mers; this halves the index (Section 6). Our streaming search for this index, `runs`, exploits how rarely the orientation changes along a read.
6. We adapt the fixed block correction sets of Diseth and Puglisi [15] to batched search, laying them out so that a rank query usually reads one cache line whose address follows from the query position alone and can be prefetched in one stage (Section 7). This brings the index to about 3 bits per canonical *k*-mer.

We evaluate all methods against SSHash on the collections of its authors’ benchmark and on two collections 3–15 times larger (Section 8). Together, our indexes form a new Pareto frontier for the space–time trade-off of *k*-mer lookups.

## 2 Preliminaries

A string *T* = *T* [0*..n* − 1] of length |*T* | = *n* is a sequence of symbols from the DNA alphabet Σ = {`A`, `C`, `G`, `T`}. The substring from position *i* to position *j* inclusive is *T* [*i..j*]; *T* [*i..j*) excludes *T* [*j*], *T* [*i..*] is the suffix starting at position *i*, and *ɛ* is the empty string. The same notation applies to arrays and bitvectors. A *k*-mer is a string of length *k*. The *colexicographic* (colex) order of strings is the lexicographic order of their reverses.

**Definition 1** *(k-spectrum)* The *k*-spectrum *S_k_*(*T*) of a string *T* is the set of distinct *k*-mers of $*^k^T*, where $ ∈*/* Σ is smaller than every symbol of Σ. The *k*-spectrum of a set of strings is the union of their *k*-spectra.

The *k*-mers containing $ are called *dummy k*-mers. They ensure that every prefix of every *k*-mer is a suffix of some *k*-mer of the spectrum, so that a search can proceed one character at a time. A *k-mer lookup* for a string *P* of length *k* returns the colex rank of *P* in *S_k_*(*T*), or ⊥ if *P* ∈*/ S_k_*(*T*).

The *reverse complement* of a DNA string is the string reversed, with each base replaced by its complement (`A` ↔ `T`, `C` ↔ `G`); a *k*-mer and its reverse complement describe the same sequence read from opposite strands. The *canonical* form of a *k*-mer is the smaller of the *k*-mer and its reverse complement. In the node-centric *de Bruijn graph* of a *k*-spectrum, the nodes are the *k*-mers, with an edge from *x* to *y* whenever *x*[1..] = *y*[0*..k*−1). A *unitig* is a maximal path in which every node except the first has in-degree one and every node except the last has out-degree one.

The spectral Burrows–Wheeler transform (SBWT) [2] represents a *k*-spectrum as a sequence of subsets of Σ; we refer to [2, 1, 4] for a more comprehensive treatment.

**Definition 2** *(SBWT)* Let *A*_1_*, … , A_|Sk |_* be the elements of a *k*-spectrum *S_k_* in colex order. The SBWT is the sequence of sets *X*_1_*, … , X_|Sk |_* with *X_i_* ⊆ Σ such that *X_i_* = ∅ if *i >* 1 and *A_i_*[1*..k* − 1] = *A_i−_*_1_[1*..k* − 1], and otherwise *X_i_*= {*c* ∈ Σ | *A_i_*[1*..k* − 1]*c* ∈ *S_k_*}.

In other words, *X_i_* holds the labels of the outgoing edges of *A_i_* in the de Bruijn graph, recorded only for the first *k*-mer of each *suffix group*, a maximal run of *k*-mers sharing the same suffix of length *k* − 1. Figure S1 in the appendix shows the *k*-mers and the SBWT of *T* = ACATACACGTCAGCACGA for *k* = 3.

**Definition 3** *(Subset rank)* Given a sequence *X*_1_*, … , X_n_* of subsets of Σ, a subset rank query `subset-rank`(*i, c*) returns the number of subsets *X_j_* with *j < i* such that *c* ∈ *X_j_*.

*Lookup.* The *colex interval* [*s, e*]*_α_* of a string *α* holds the colex ranks of the *k*-mers of *S_k_* that have *α* as a suffix. Given [*s, e*]*_α_* and *c* ∈ Σ, *ExtendRight*([*s, e*]*_α_, c*) returns [*s′, e′*]*_αc_*, or ⊥ if it is empty, using two subset rank queries [2]: *s′* = *C*[*c*] + `subset-rank`(*s, c*) + 2 and *e′* = *C*[*c*] + `subset-rank`(*e* + 1*, c*) + 1, with ⊥ if *s′ > e′*, where *C*[*c*] is the number of occurrences in the SBWT of characters smaller than *c*. A lookup of *P* starts from [1, |*S_k_*|]*_ɛ_* and applies *ExtendRight* once per character of *P*. In this example, looking up `ACG` visits [1, 16]*_ɛ_* → [2, 7]_A_ → [8, 10]_AC_ → [14, 14]_ACG_, so its colex rank is 14. In practice, a lookup starts from a precomputed *prefix lookup table* holding the intervals of all *p*-mers, which saves its first *p* steps; we use *p* = 8.

*Streaming search.* A streaming query is a string longer than *k*, typically a sequencing read; we call it a *read*. We call the *k*-mer starting at position *i* of a read its *window i*. A streaming search reuses the interval of one window for the next, and needs one of two adjuncts to the SBWT.

**Definition 4** *(LCS array, suffix-group bitvector)* Let *A*_1_*, … , A_|Sk |_* be the elements of *S_k_* in colex order. The *longest common suffix* (LCS) array has LCS[1] = 0 and, for *i >* 1, LCS[*i*] equal to the length of the (possibly empty) longest common suffix of *A_i_* and *A_i−_*_1_. The *suffix-group bitvector V* has *V* [*i*] = 1 iff LCS[*i*] = *k* − 1.

If *ExtendRight*([*s, e*]*_α_, c*) fails, the LCS array gives the interval of *α*[1..], dropping the first character, and the search continues from there without restarting [1]. The suffix-group bitvector supports this only for |*α*| = *k*, that is, from a *k*-mer to its suffix group; otherwise the next window is searched *from scratch*, which we call a *restart*. We call a step that extends the interval of the previous window a *continuation*. The LCS array costs ⌈log *k*⌉ bits per *k*-mer and the bitvector one bit, a space–time trade-off.

*Matrix representation.* Subset rank queries on the SBWT can be supported in many ways. We mostly use a binary matrix *M* with one row per *c* ∈ Σ, such that *M* [*c*][*j*] = 1 iff *X_j_* contains *c* (Figure S1). Then `subset-rank`(*i, c*) = rank_1_(*M* [*c*]*, i*), where rank_1_(*B, i*) is the number of 1-bits in *B*[0*..i*), answered by a rank structure on each row. With rank9 [16, 34], this is the fastest subset rank structure in practice [2], at 5 bits per subset: 4 for the matrix column, plus the overhead of the rank structures.

## 3 Speeding up *k*-mer lookups by mini-batching queries

A *k*-mer lookup in the SBWT gives rise to a chain of memory latencies. In particular, each search step depends on the interval computed in the previous step, and each step accesses essentially random locations in the subset rank structure [4]. In our profiling of the matrix with rank9, we measure on average 1.7–2.3 last-level cache (LLC)^1^ misses per search step, each step taking 95–98 ns—roughly the time it takes the CPU to resolve one cache miss. The misses of a step’s two rank queries presumably overlap. A core can serve about *F* = 12 cache misses at the same time (its L1 fill buffers), of which a single query uses about two.

We now show that this unused capacity can be largely utilized by processing a small batch of queries—a *mini-batch*—in a loop, effectively increasing the amount of independent work for the CPU to perform while waiting for memory. A similar line of enquiry was recently explored by Groot Koerkamp in the context of the FM-index [17], though the idea has precursors in work on BWT inversion by Kärkkäinen, Kempa and Puglisi [21] and in Movi [36, 37], which processes several reads concurrently with prefetching. We build on these ideas and introduce variations of our own.

Because each search step already overlaps two cache misses, processing many queries at once can achieve a speedup of at most *F/*2 times. Searching queries a small batch at a time in lockstep, we measure a speedup of only 1.6 times, almost all of it already with batches of size two. This is because one step is over 100 instructions, and the CPU’s out-of-order look-ahead window (224 micro-ops on our CPU) covers instructions for only the next one or two queries. This implies batching alone can only go so far.

In a similar vein to Groot Koerkamp [17], we obtain further gains in performance via *memory prefetching*. In particular, as soon as a step of query *q_j_* is done—its interval extended by one character by answering its endpoint subset rank queries—we issue a prefetch of the cache lines the *next* step will read and move on to the next item in the batch (*q_j_*_+1_); on returning to *q_j_* one sweep over the batch later, these lines reside in cache. This lifts the gain from 1.6 to 4.3–5.0 times. Profiling shows batches of size 16–64 to work best. Finally, we keep the batches full by *refilling* a slot as soon as the search for its *k*-mer ends.

### 3.1 A rank structure hitting one cache line

With the latency hidden, what remains is to try to reduce the number of cache lines involved in each search step. Rank9 [34, 16], as implemented in the simple-sds crate we used in our experiments above, reads two cache lines per rank query: its rank support structure (an absolute count and seven cumulative 9-bit counts per 512-bit block) is in an array separate from the data bits making up the actual bitvector itself.

We now describe a novel *one-line rank*, in which each rank query reads exactly one aligned 64-byte cache line. One-line rank can be thought of as an in-line version of rank9: like SPIDER [22] and BiRank [17], it stores the counts inside the cache line of the data, and, as in rank9, the counts are an absolute count and cumulative per-word counts, so that a rank query needs no separate table access and exactly one popcount.^2^ The closest published structure to one-line rank is BiRankR9, one of the variants proposed by Groot Koerkamp [17], with rank9’s field widths: a 64-bit absolute count and six 9-bit counts, 128 header bits and 384 data bits, or 33.3% overhead. One-line rank narrows each field to what it needs (a 40-bit absolute count and cumulative counts of 6–9 bits) fitting all fields into 96 bits, so that a line holds 416 data bits, for 23.1% overhead, similar to rank9’s. Figure 1 shows the layout of a line. No field crosses a word boundary.

**Figure 1:**
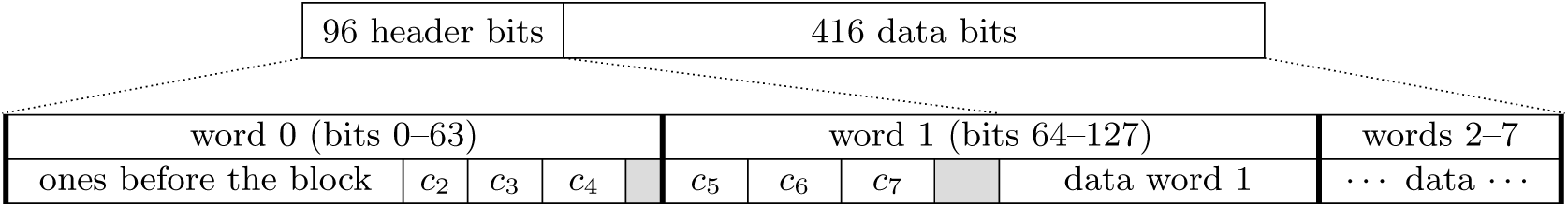
A line of one-line rank. Top: the 512-bit line, 96 header bits and 416 data bits. Bottom: the header zoomed in, to scale, with the data words 2–7 collapsed (grey: unused bits; thick lines: word boundaries); *c_w_* is the number of ones before data word *w* within the line.

We answer rank(*i*) for bit *i* of a row of the matrix with one-line rank as follows. The required line has index ⌊*i/*416⌋, the offset within it is *p* = 96 + (*i* mod 416), and the word *w* = ⌊*p/*64⌋. The rank answer then is the absolute count, plus the cumulative count of word *w* (0 for *w* = 1), plus the popcount of word *w* masked to the bits before *p* (and, for *w* = 1, to its data half). From here on, all matrix rows use this layout.

We conclude this section with one small improvement. When the endpoints of a search interval fall within the same line, the two rank queries involved can share work. In particular, we rank both ends of the interval using one specialized (*rank pair*) function, which accesses the appropriate rank structure once. Over the single-query search with rank9, the cumulative speedup of *k*-mer lookup is 1.6 times with batching alone, 4.3–5.0 times with prefetching, 4.4–5.4 times with refilling, 7.0–7.4 times with the one-line rank, and 7.1–7.5 times with the rank pair.

## 4 Mini-batching in streaming mode

Realizing the gains of batching and prefetching in streaming search (Section 2) is harder, because progress through the query reads of a batch can become uneven and the actions needed to progress in different queries diverge. This happens when a mismatch occurs at a given position in some queries, but not in others. Most positions need a single interval extension, but, if an extension fails, the matching process must back off using the streaming support structure. In this way, processing a position across a batch may involve several dependent steps of different kinds, and the search can become full of data-dependent branches. Moreover, the two types of streaming search (LCS-based and bitvector-based; Section 2) diverge in different ways. With the LCS array, a query backs off one character at a time, for one or more rounds, each requiring a scan of the LCS array for an unknown distance. With only the suffix-group bitvector, a query can back off only from a full *k*-mer to its suffix group, and otherwise must restart. This search thus mixes continuations of previously found *k*-mers, which take one step, with restarts, which take up to *k* steps; interleaved in one loop, they stall each other. We therefore design different batching approaches for LCS-based streaming and bitvector-based streaming, tailored to how query processing diverges in each scheme.

We batch the streaming search with the LCS array as follows. A mini-batch holds 32 query reads, and each sweep over the batch does a single step per query. After an extension we prefetch what the query needs next, which is either the rank data of the next extension if it succeeded, or the LCS entries at both interval ends if it failed. In the next sweep, the queries that failed are processed as one group. Each backs off with the prefetched LCS entries and extends again; if that fails, it continues to back off and extend until an extension succeeds. In these continued back-off steps, the next LCS entries lie next to those just loaded. Similarly, during the back-offs the rank data for the new intervals usually fall in cache lines already loaded. Once an extension succeeds, the query prefetches the rank data of its next extension and rejoins the group of extending queries.

With the suffix-group bitvector, we process continuations and restarts in separate loops. A batch contains up to 64 query reads and when a read is processed to its end, it is replaced with a new read from the query set. In a given round of the batch loop, every read whose last *k*-mer was found is extended by its next character, using rank data prefetched in the previous round. Because the *k*-mer was found, this takes a single rank query. This extension fails either if the next *k*-mer is absent, or if the outgoing edge is stored at another *k*-mer of the same suffix group (Section 2). If it fails, we prefetch the needed region of the suffix-group bitvector and continue along the batch. When next returning to this read, we issue two rank queries delineated by the suffix group. If this extension fails too, the read must restart and is placed in a *restart queue*. Whenever the restart queue fills to 32 reads, or no read can continue, their *k*-mers are searched together with the batched *k*-mer search of Section 3.

On reads without substitutions, whose *k*-mers are all in the index, batching speeds up both streaming searches considerably (Section 8.2), though by less than it speeds up *simple k*-mer lookups (Section 3). These speedups are eroded as substitutions increase: gradually with the LCS array, and quickly with the bitvector, where every substitution causes up to *k* restarts. Most of these restarts are redundant, however, as we show next: a failed search for one *k*-mer can be used to infer several absent *k*-mers, which can then be skipped.

## 5 Skipping on mismatch: Removing the LCS array

When the search of window *w* fails after its first *l* characters, those *l* characters occur in no *k*-mer of the index at all: by the dummy *k*-mers, any string that occurs inside a *k*-mer of the index is a suffix of some *k*-mer, and so has a non-empty interval (Section 2). Hence, every window of the read that contains them is absent too: the windows starting at *w* + *l* − *k, … , w*, that is, *k* − *l* + 1 windows, can be decided by one search (Figure 2). Importantly, only windows to the *left* of the failed one can be skipped, while the streaming SBWT-search proceeds through a read left to right: by the time a window fails, the windows it could be used to eliminate have already been searched.

**Figure 2:**
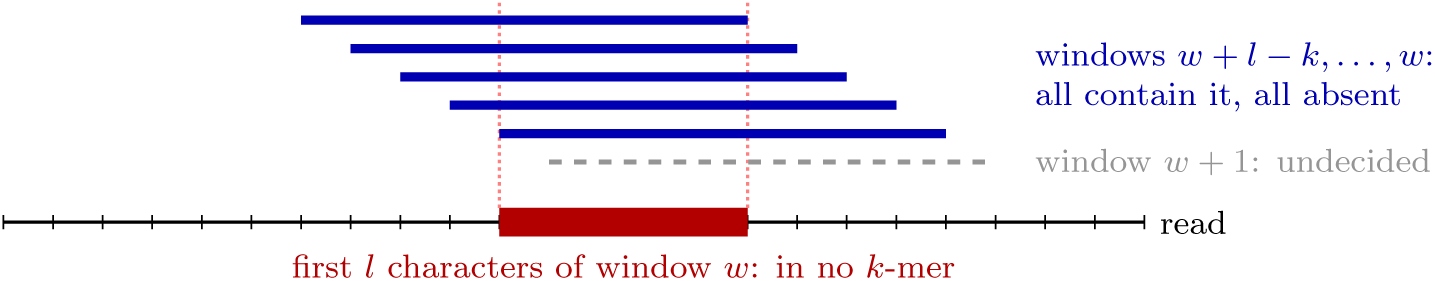
The search of window *w* fails after *l* characters (here *k* = 9, *l* = 5). Every window that contains these *l* characters is absent: *k* − *l* + 1 windows, decided by one search.

However, we can exploit the observation in streaming search as follows. The continuations still go from left to right; after a window that is not found, the search *looks ahead* to a window whose failure reaches back, and fills the remaining gap from right to left, each failure there skipping further left. We call this method `forward-skip`. There are two cases to consider: a failure at a continuation and a failure after a restart. The cases are explained in Table 1.

**Table 1:**
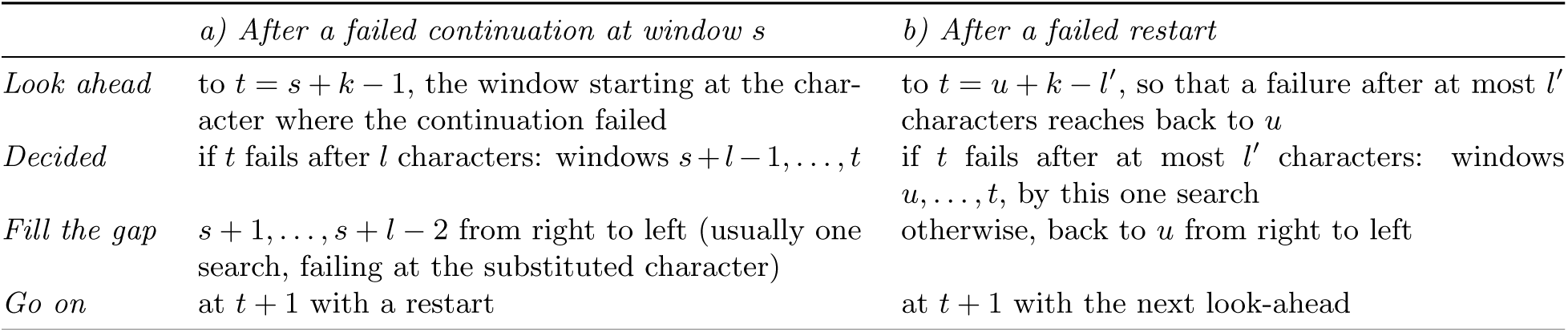
The two cases of `forward-skip`; *u*: the first undecided window, *l′*: the length after which the previous search failed (initially 15). The value of *l′* affects only speed, as any gap left is filled from right to left. If *t* is found, the gap is filled the same way and the read continues from *t*.

|  | *a) After a failed continuation at window s* | *b) After a failed restart* |
|---|---|---|
| *Look ahead* | to t = s + k − 1, the window starting at the char-acter where the continuation failed | to t = u + k − l′, so that a failure after at most l′ characters reaches back to u |
| *Decided* | if t fails after l characters: windows s + l − 1, …, t | if t fails after at most l′ characters: windows u, …, t, by this one search |
| *Fill the gap* | s + 1, …, s + l − 2 from right to left (usually one search, failing at the substituted character) | otherwise, back to u from right to left |
| *Go on* | at t + 1 with a restart | at t + 1 with the next look-ahead |

All restarts, including the look-aheads and the searches that fill gaps, go into a single restart queue, and continuations are batched as in Section 4. Whenever the restart queue holds 32 reads, their restarts are processed together with the batched *k*-mer search of Section 3. With skipping, the bitvector search keeps pace with the LCS array search and overtakes it as substitutions increase, at 60% of its memory (Section 8.2).

## 6 One-orientation SBWT indexing

With the LCS array no longer required, what remains to shrink is the SBWT itself. Indexes such as Themisto [3] store every *k*-mer together with its reverse complement, so that a read need only be searched in one orientation. Keeping one orientation of each *k*-mer halves the index, but a read must then be searched in both. Similar ideas appear in the primary unitigs mode of Metagraph [20] and in the context of finimizers [5]; here we focus on choosing orientations that minimize dummy *k*-mers, and on searching the index efficiently.

The orientations must be chosen carefully because every source node of the de Bruijn graph of the selected *k*-mers, that is, every *k*-mer with no predecessor among them, requires up to *k* dummy *k*-mers to support it. We choose orientations for whole unitigs, working on the compacted de Bruijn graph, where each node is a unitig and there is an edge from unitig *v* to unitig *u* if the last (*k* − 1)-mer of *v* is equal to the first (*k* − 1)-mer of *u*. We denote the reverse complement of unitig *v* with `rc`(*v*). Since our starting SBWT is closed under reverse complementation, both *v* and `rc`(*v*) always exist as unitigs of the graph. The problem is now to select one unitig from each pair {*v,* `rc`(*v*)} (one choice if *v* = `rc`(*v*)), so that the induced subgraph of the selected unitigs has as few source nodes as possible.

We use a greedy heuristic, with no claim of optimality: until all unitigs are visited, we select an unvisited unitig *v* and run a breadth-first search over unvisited unitigs from *v*, first forwards and then backwards, selecting each unitig *u* it reaches; each time, both *u* and `rc`(*u*) are marked visited. Each unitig is thus selected in the orientation in which an already selected unitig reaches it, so that it has a predecessor and is not a source; the backward search does the same for *v* itself.

In our experiments, with this heuristic, the number of dummy *k*-mers grows from at most 0.5% on the two-orientation indexes to 1.6–2.9% on the one-orientation indexes, while the number of *k*-mers is halved: the one-orientation SBWT takes 50.7–51.5% of the space of the two-orientation one.

What remains is to search it efficiently, as a read may have to be searched in both orientations. Searching each read twice, once in each orientation, is wasteful: windows found in one orientation incur mismatches in the other, creating the worst case for the streaming search. But a read from a genome related to the indexed ones follows a unitig for many windows before it may switch to another. Since the *k*-mers of the index come from whole unitigs, consecutive windows within a unitig keep their orientation; and since the heuristic above selects each unitig in the orientation in which a neighbouring selected unitig reaches it, a read that crosses from one unitig into the next usually keeps its orientation too. Therefore the orientation rarely changes along a read: windows are found in long runs, either as they are or as their reverse complements.

We call our streaming search on the one-orientation SBWT, which uses the suffix-group bitvector, `runs`. It first follows a run from each end of the read, as in cases a) and b) below, and then decides the windows between the two runs as in cases c) and d). Figure 3 shows the four cases:

*a) A forward run* (blue, left in Figure 3): from the first window of the read, the search follows the windows left to right with continuations, until a window *f* is not found.
*b) A reverse run from the end* (orange, right), once per read: windows found only as reverse complements can be followed with continuations only from right to left, as the SBWT extends to the right. So the reverse complement of the read is searched from the read’s last window, until a window is not found.
*c) A reverse run in the middle* (green and orange): if the forward run stops at window *f* (possibly *f* = 0) and the reverse complement of window *f* is in the index, a run of reverse complements starts at *f* , which can only be followed from its right end *r*. We find *r* by galloping: the reverse complements of windows *f* + 1*, f* + 3*, f* + 7*, …* are searched by restarts until one is not found, and bisection between the last window found and the first not found then gives *r*. The run is then followed back from *r* to *f* , as in b), and the forward run goes on after *r*. A gap in the run only costs extra searches: every window is still decided.
*d) Windows in neither orientation* (red): a window *g* found in neither orientation is decided by skips (Section 5), one per orientation. The search of the reverse complement at *g* fails after *l* characters, which decides the reverse complements of windows *g, … , g* +*k* −*l*; one search of window *g* +*k* −*l*, skipping leftwards, then usually decides their forward *k*-mers. A stretch absent in both orientations thus costs two searches per skip, against one on the two-orientation index.

**Figure 3:**
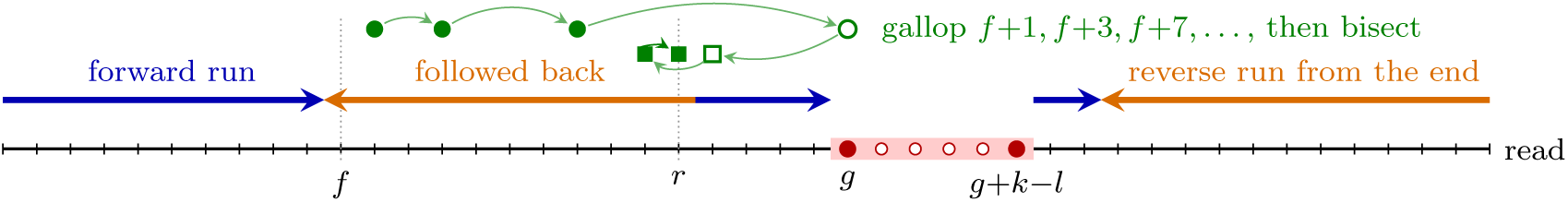
The `runs` search on a read against a one-orientation SBWT. Blue: runs found in the read’s orientation, followed left to right; orange: runs found as reverse complements, followed right to left; green: galloping (circles) and bisection (squares) for the end *r* of a reverse run starting at *f* (filled: found, open: not found); red: windows in neither orientation, decided by a skip (filled: searched, open: skipped).

Once a window is found again, the search goes on as in a) or c).

Every window found is found once, by a continuation or a restart. We batch `runs` as `forward-skip` (Section 5): a batch consists of 64 reads, with 32 restarts at a time. At half the space, it takes 1.3–2.1 times the time of `forward-skip` on the two-orientation index (Section 8.2).

## 7 Correction sets for subset rank: a final squeeze

Having eliminated the LCS array (Section 5) and halved the number of *k*-mers indexed (Section 6), the remaining option to reduce space further is to reduce the size of the subset rank data structure used to traverse the SBWT. To this end, we now tailor the recent *correction-sets* subset rank structures [15] to batched streaming queries. These structures use around 3 bits per subset at the cost of slower queries. This matters little to a single query, bound by memory latency, but a lot to our batched searches, whose performance is achieved by hiding memory latency and so is bound by instructions. We now describe a variant of the *fixed block* correction-sets (FBCS) [15], designed with mini-batches in mind.

We store the subset sequence as a string of *base symbols* plus a few *corrections*. The base symbol of a subset is its smallest character, or A if it is empty; as most subsets are singletons, it usually describes the subset exactly. Corrections record where it does not: for A, the positions of empty subsets; for C, G and T, the positions of subsets that contain the character but have a smaller base symbol. Then `subset-rank`(*i, c*) is the number of base symbols *c* before position *i*, minus the corrections of A before *i* if *c* = `A`, and plus the corrections of *c* before *i* otherwise. For example, {`A`}, ∅, {`C`, `G`} have base symbols A, A, C, a correction of A at position 2 and one of G at position 3.

The subsets are divided into blocks of 184, each stored as one aligned 64-byte line, so that the block of subset *i* is line ⌊*i/*184⌋. The symbol counts before each block are kept in three levels (Figure 4): 64-bit absolute counts every 2^17^ upper blocks, 32-bit counts every upper block of 178 blocks (32 752 subsets), and 15-bit counts in the block itself; the upper level is small enough to stay in the last-level cache.

**Figure 4:**
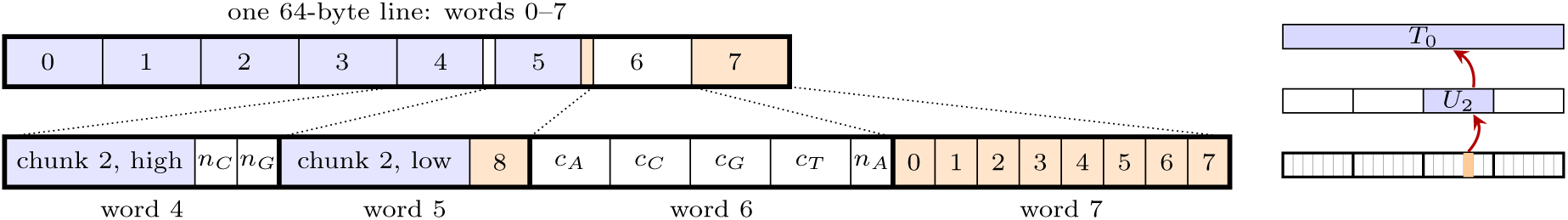
Left: an FBCS block, its eight words above, and words 4–7 zoomed in below. The base symbols of its 184 subsets (blue) fill most of words 0–5; *c_A_, … , c_T_* are the counts before the block, *n_A_*, *n_C_*, *n_G_* the numbers of its corrections of A, C and G, and 0–8 the corrections themselves (orange), one byte each. An overflowing block uses word 7 to store a pointer to the overflowing data. Right: the counts before a position in three levels (not to scale): in the block, in the upper array (*U*), and in the super level (*T*). A rank query in the block of position *i* (orange) adds one entry of each level (red arrows).

Besides the counts, a block holds the base symbols of its 184 subsets and up to 9 corrections, each stored as its position within the block, so that a rank query at position *i* counts the corrections of its character below *i* mod 184. A block with more than 9 corrections *overflows*: its extra corrections go to a separate array, to which the block keeps a pointer. Up to 35% of the blocks overflow, depending on the collection (Table S6), and up to 3*/*4 of the rank queries that land in an overflowing block need the overflow array.

*Prefetching.* Batching hides memory latency by prefetching, one step ahead, the data that each query’s next rank query will read. For the one-line rank of Section 3.1, this is a single cache line whose address follows from the query position alone. The fixed block layout is designed to come as close to this as possible. The block of position *i*, too, follows from *i* alone, as does the word of upper block counts it needs, so the prefetch has a single stage: the block and its upper count are requested together, with no dependent read. Only a block that overflows needs a further, dependent read of its overflowing corrections. Eight prefetch calls later, by which time the block has arrived, the prefetch reads the block’s overflow pointer and, if it is set, prefetches the overflowing corrections as well.

What remains different from the bit matrix is computation: a rank query on the FBCS costs about 120–175 instructions against about 45, and once the memory latency is hidden, batching cannot remove this cost. We batch the FBCS streaming search as we do the matrix search, with `runs`. Since each rank query might need to read its data from more than just one cache line, we use a smaller batch size: 32 instead of 64. At 2.8–3.3 bits per canonical *k*-mer for the index alone (versus 5.1), the FBCS trades speed for space, a new point on the Pareto front (Section 8.2). We consider the FBCS only on the one-orientation index: with both orientations it would take about 5.7 bits, more than the faster one-orientation bit matrix.

## 8 Experiments

We evaluate our methods in two parts: Section 8.2 measures what each technique gains over the previous SBWT search, and Section 8.3 compares our indexes with SSHash, the leading hashing-based index.

### 8.1 Setup

All our methods are implemented in Rust (Appendix B.6); SSHash [30] is queried through a small C++ driver on the same queries. Every experiment runs on one core of an Intel Xeon Gold 6248 (2.5 GHz, 27.5 MB last-level cache). Space is given in bits per canonical *k*-mer of the collection, and time in ns per *k*-mer of the queries; we report the best of three runs.

#### Datasets

Table S1 in Appendix B.2 lists the collections and their sizes. Three test collections (E. coli, Salmonella and 60 human genomes) measure the improvements to the SBWT search, and the six collections of the SSHash benchmark [32]^3^ the comparison with SSHash. HumGut (2025)^4^ and ParaRef, 3–15 times larger than any collection of the benchmark, are used in both. For each collection and *k*, all indexes, SSHash’s and ours, hold exactly the same *k*-mers (Appendix B.4).

#### Queries

*k*-mer lookups use 10^6^ positive *k*-mers randomly sampled from the index and 10^6^ random *k*-mers. On our three test indexes and on HumGut and ParaRef, the streaming queries are 2 · 10^4^ reads of 200 bases cut from the genomes, with 0–5% random substitutions, and 10^4^ random reads of 200 bases. In the comparison with SSHash, we use three sets of real reads for each dataset, of up to 10^6^ reads each: high, reads mostly from the collection; related, reads of a related organism or sample; and low, reads mostly absent from it (Appendix B.3).

#### Indexes compared

We compare SSHash, which has its streaming support built in, with three SBWT indexes: the two-orientation matrix, searched with `forward-skip` (12 bits per canonical *k*-mer in all), the one-orientation matrix (6.1 bits) and one-orientation FBCS (about 4 bits), both searched with `runs`. The streaming support for all indexes is the suffix-group bitvector. In SBWT performance evaluation (Section 8.2) the baseline is the single-query search on the two-orientation matrix with the one-line rank and the LCS array.

### 8.2 Performance of SBWT variants

#### *k*-mer lookups

On the three test collections, a positive *k*-mer takes 1869–2043 ns with the baseline and 295–320 ns batched (Table S4), 6.3–6.5 times faster (less than in Section 3, as the baseline already uses the one-line rank); on random *k*-mers the batched search is 5.3 times faster. Profiling shows that the batched search is no longer bound by memory, but by instructions: on the bit matrix it runs at 2.5–2.7 instructions per cycle with only 0.1–0.2 last-level cache misses per *k*-mer.

#### Streaming search

Figure 5 shows the batched streaming searches of Sections 4–7 against the baseline, by substitution rate, on the three test collections, HumGut and ParaRef. With the LCS array, batching makes streaming search 4.7–5.2 times faster than the baseline on reads without substitutions. The gain shrinks to 2.4–2.7 times at 5% substitutions and about 2 times on random reads, as matches get shorter and the search backs off more often. Replacing the LCS array by the suffix-group bitvector with `forward-skip` cuts the space from 20 to 12 bits per canonical *k*-mer at little or no cost in time: up to 2% substitutions the two searches are within 10% of each other, and at 5% `forward-skip` is faster on every collection, by 1.2–1.6 times. On random reads, where skipping decides most windows without a search, it is 3–8 times faster. The one-orientation index with `runs` halves the space again, to 6.1 bits, at 1.3–2.1 times the time of `forward-skip`; it is still 1.7–3.2 times faster than the baseline on reads with up to 5% substitutions, and 4.7–9.6 times faster on random reads. FBCS cuts the space to about 4 bits, at 2.2–3.6 times the time of the one-orientation matrix. At a fifth of the baseline’s space, it runs at 0.5–1.2 times its speed on reads with substitutions and 1.7–3.9 times faster on random reads.

**Figure 5:**
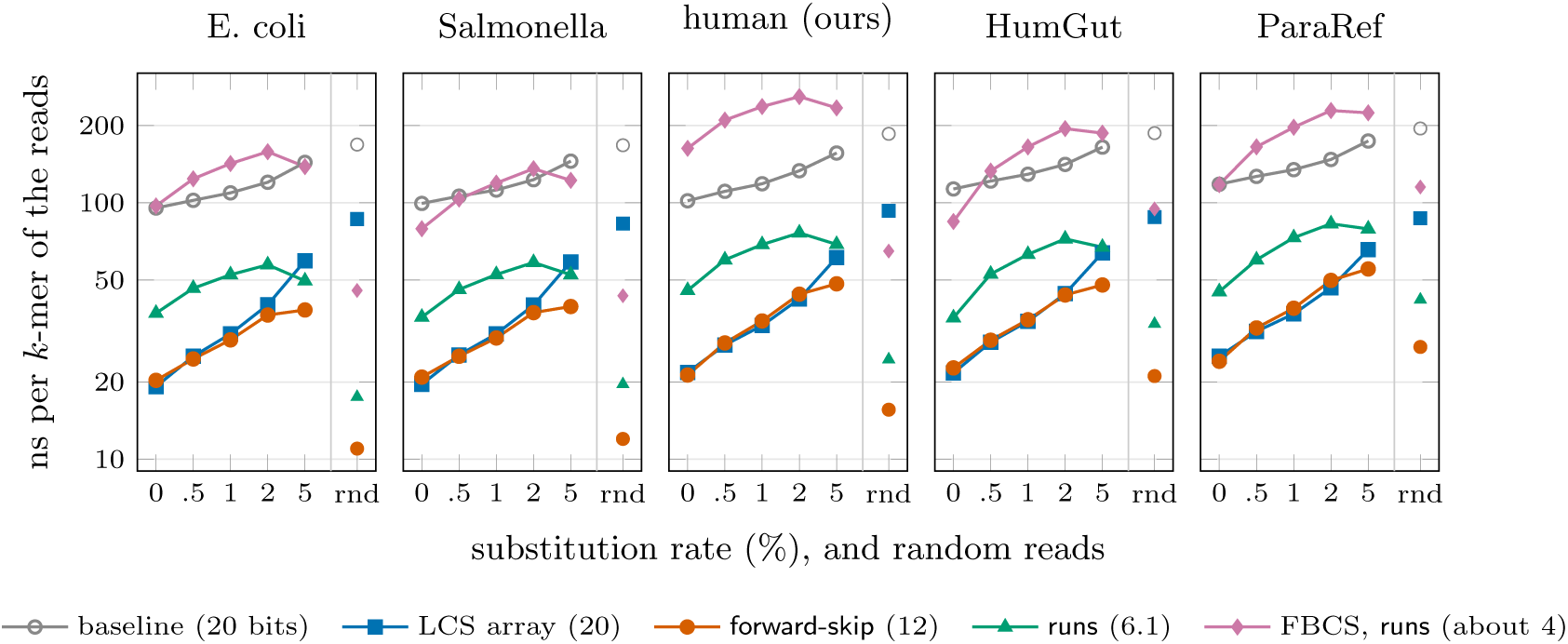
Streaming queries by substitution rate on the test indexes, HumGut and ParaRef (*k* = 31): ns per *k*-mer of the reads, logarithmic scale; reads of 200 bases with 0–5% random substitutions, and random reads (rnd). In brackets, bits per canonical *k*-mer of index and streaming support. All values in Table S5.

### 8.3 Comparison with SSHash

SSHash, a minimizer-based hash index, is the reference *k*-mer dictionary: in its authors’ benchmark it is the fastest for all queries, ahead of the single-query SBWT. We compare our three indexes with it on the six collections of its benchmark and on HumGut and ParaRef, at *k* = 31 and *k* = 63. Figure 6 shows the speedup of our streaming searches over SSHash on every read set, together with the space of each index, and Table 2 summarizes the *k*-mer lookups (all values: Tables S7 and S8). SSHash answers queries one at a time, as it is used in practice; whether it would gain as much from batching is an open question (Section 9).

**Figure 6:**
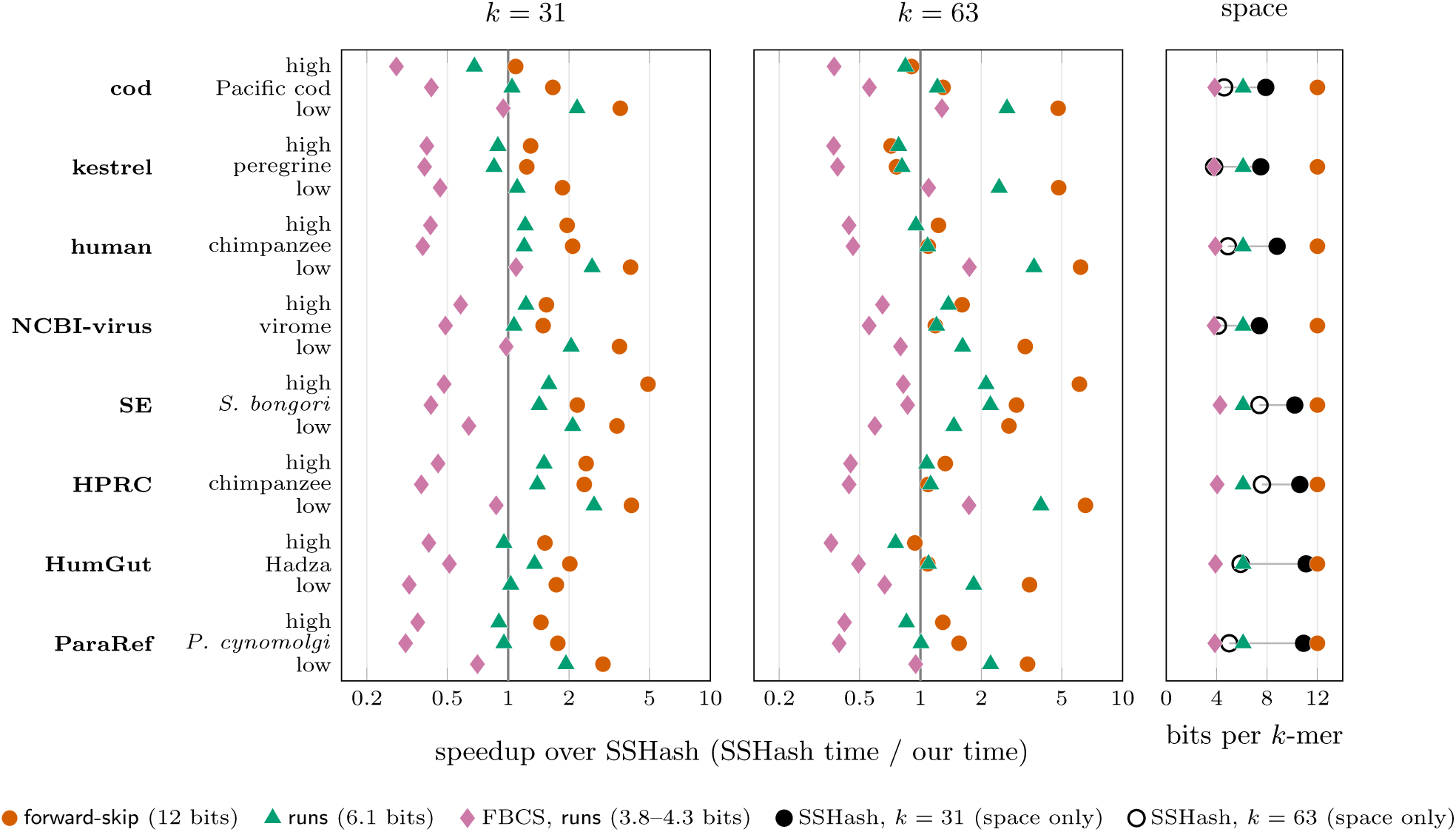
Read queries: speedup of our batched streaming searches over SSHash on every collection and read set (related reads named by organism or sample), logarithmic scale. Right of the vertical line, our search is faster. In brackets, bits per canonical *k*-mer of index and streaming support (SSHash: 7.4–11.1 bits at *k* = 31, 3.8–7.6 at *k* = 63). Right: bits per canonical *k*-mer of each index with its streaming support; SSHash at *k* = 31 (filled) and *k* = 63 (open), joined by a line; our indexes take the same space at both *k*, except FBCS, whose size differs by at most 0.14 bits (shown: *k* = 31). All values in Table S7.

**Table 2:** *k*-mer lookups: median time over the eight collections (the six of the SSHash benchmark, HumGut and ParaRef), in ns per *k*-mer, with the range in brackets; 10^6^ positive and 10^6^ random *k*-mers; bits per canonical *k*-mer of the index alone. All values in Table S8.

| index | *k* = 31 |  |  | *k* = 63 |  |  |
|---|---|---|---|---|---|---|
|  | bits | positive | random | bits | positive | random |
| SSHash | 7.4–11.1 | 550 (438–628) | 359 (312–444) | 3.8–7.6 | 791 (630–1002) | 528 (482–603) |
| two-or. matrix | 10 | 308 (301–392) | 132 (122–212) | 10 | 724 (708–920) | 166 (155–253) |
| one-or. matrix | 5.1 | 363 (338–451) | 230 (207–365) | 5.1 | 800 (734–938) | 267 (243–405) |
| one-or. FBCS | 2.8–3.3 | 1289 (856–1543) | 793 (543–1263) | 2.8–3.1 | 2164 (1755–2456) | 709 (588–1137) |

#### Reads

At *k* = 31, `forward-skip` on the two-orientation index is faster than SSHash on every read set, by 1.1–4.9 times, at 12 bits per canonical *k*-mer against SSHash’s 7.4–11.1. At *k* = 63 it is faster on 20 of the 24 read sets; on the other four (the high reads of cod, kestrel and HumGut, and the related reads of kestrel) SSHash is up to 1.4 times faster. On reads mostly absent from the collection, the gap is consistently large, 1.7–4.1 times at *k* = 31 and 2.7–6.6 times at *k* = 63, as `forward-skip` decides most windows without a search. The one-orientation index with `runs`, at 6.1 bits, is smaller than SSHash on every collection at *k* = 31 and still faster than SSHash on 18 of the 24 read sets, at both *k*. FBCS, at 3.8–4.3 bits, trades speed for space: it is slower than SSHash on all but one read set at *k* = 31, by up to 3.6 times, and on all but four at *k* = 63.

#### Lookups

At *k* = 31, the two-orientation matrix is faster than SSHash on every collection, 1.5–1.9 times on positive *k*-mers and 2.1–2.7 times on random ones. At *k* = 63 it is still 2.4–3.2 times faster on random *k*-mers, but on positive *k*-mers SSHash is faster on four of the eight collections, by up to 1.12 times. The one-orientation matrix, at half the space, is at *k* = 31 faster than SSHash on every collection, by 1.2–1.6 times, on both positive and random *k*-mers. FBCS takes 1.2–3.0 times SSHash’s time on all *k*-mers.

#### Space

The size of an SBWT is set by its number of *k*-mers, not by *k* or by how the *k*-mers are arranged, so our matrix indexes take the same space per *k*-mer on every collection, and FBCS nearly so (Figure 6, right). The size of SSHash depends on how the data falls into super-*k*-mers: 7.4–11.1 bits per *k*-mer at *k* = 31 and 3.8–7.6 bits at *k* = 63, where it is sometimes smaller than our one-orientation matrix index, and on kestrel as small as FBCS. The space of our indexes is thus known in advance, from the number of *k*-mers alone.

## 9 Concluding Remarks

We have shown that *k*-mer lookup with the SBWT can be made much faster and smaller by engineering aimed at this data structure: mini-batching with prefetching, a one-line rank structure, skipping on mismatch, one-orientation indexing, and a compact subset rank structure adapted to batched search. The resulting indexes answer streaming queries faster than SSHash at comparable space, or use about 4 bits per canonical *k*-mer; and as an SBWT’s size is set almost entirely by its *k*-mer count, its memory needs are known in advance.

On HumGut, for example, our smallest index takes about 7 GB with its streaming support, against about 20 GB for SSHash, and SBWT-based tools such as the pseudoaligner Themisto [3] can use them directly.

Once memory latency is hidden, the bottleneck moves from cache misses to instructions (Section 8.2): the compact structure is slower not because it reads more memory, but because each rank query costs three to four times as many instructions. For batched search, subset rank structures should minimize computation per query, while keeping the memory each query reads predictable from its position alone.

Several directions remain open. Since batched search is bound by instructions, SIMD across the queries of a batch might narrow the gap between the correction-set structure and the bit matrix. Our experiments use one core; each thread could run its own mini-batches, but whether the gains hold when many cores share memory bandwidth remains to be measured. Finally, mini-batching is likely to speed up SSHash and possibly other *k*-mer indexes too (e.g., [33]), though the details will differ. We leave this for others to explore.

## A SBWT matrix example

Figure S1 shows the matrix representation (Section 2) of the SBWT for the running example of Section 2, *T* = ACATACACGTCAGCACGA with *k* = 3.

**Figure S1:**
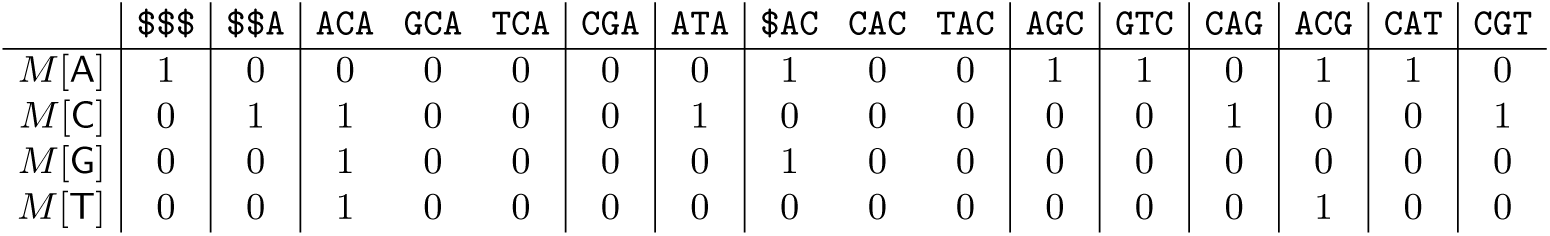
Matrix representation of the SBWT for ACATACACGTCAGCACGA with *k* = 3: the *k*-mers in colex order (top) and, for each character *c*, the row *M* [*c*], whose column for a *k*-mer is 1 iff its subset contains *c*. Vertical lines separate the suffix groups, maximal runs of *k*-mers sharing the same suffix of length *k* − 1; only the first *k*-mer of each group has outgoing edges.

## B Experimental setup

### B.1 Machine and compilation

All experiments run on one core of a node with two Intel Xeon Gold 6248 CPUs (2.5 GHz, 20 cores each, 27.5 MB last-level cache); every run is pinned to one core, with the index in the memory of that core’s NUMA node, on transparent huge pages. Our code is compiled with Rust 1.96 in release mode with -C target-cpu=native. We also align branches to 32-byte boundaries,^5^ which removes code-placement effects of up to 11% on this CPU. Rank9 is that of the simple-sds crate, compiled with the same flags. SSHash and its driver are compiled with GCC 15.2 and -march=native (Section B.5). We report the best of three runs.

### B.2 Collections

Table S1 lists the collections. Our own collections are 3 682 E. coli assemblies from NCBI (ZeSide note to avoid pencil, Gradescope or the scanner seems to increase the contrast and rubbed out pencil comes through darker and can be as dark as your real answer so I think we would prefer pen that is scribbled out rather than erased pencil.nodo 10.5281/zenodo.6577997), the Salmonella genomes of the 661k collection [8], 60 human genomes from the Human Pangenome Reference Consortium^6^ (3.04 · 10^9^ canonical 31-mers, against GRCh38’s 2.51 · 10^9^), HumGut, version 2025 (31 225 human gut genomes) [18]^7^, and ParaRef (831 parasite genomes, Zenodo 10.5281/zenodo.13744644) [28]. The six collections of the SSHash benchmark^8^ are cod (*Gadus morhua*, assembly gadMor3.0), kestrel (*Falco tinnunculus*, FalTin1.0), human (GRCh38), NCBI-virus (18 836 RefSeq virus assemblies), SE (534 751 *Salmonella enterica* genomes) and HPRC (the human pangenome of the Human Pangenome Reference Consortium).

**Table S1:** The collections and their numbers of canonical *k*-mers (10^9^).

| our collections |  |  |  | SSHash benchmark |  |  |  |
|---|---|---|---|---|---|---|---|
|  |  | *k* = 31 | *k* = 63 |  |  | *k* = 31 | *k* = 63 |
| E. coli | 3 682 assemblies | 0.17 | — | cod | gadMor3.0 | 0.50 | 0.56 |
| Salmonella | 661k coll. [8] | 0.31 | — | kestrel | FalTin1.0 | 1.15 | 1.16 |
| human (ours) | 60 genomes | 3.04 | — | human | GRCh38 | 2.51 | 2.77 |
| HumGut | 31 225 gut genomes [18] | 14.2 | 19.5 | NCBI-virus | 18 836 viruses | 0.38 | 0.41 |
| ParaRef | 831 parasites [28] | 54.8 | 61.0 | SE | 534 751 *S. enterica* | 0.89 | 1.52 |
|  |  |  |  | HPRC | human pangenome | 3.72 | 5.93 |

### B.3 Read sets

For the comparison with SSHash, each collection has three read sets of up to 10^6^ reads. For the six benchmark collections, high and low are the benchmark’s own read sets (low: reads of another organism), and related is an Illumina run of a related organism from ENA (Pacific cod, peregrine falcon, chimpanzee, a viral metagenome, *S. bongori*). For HumGut and ParaRef, high is an Illumina run of a genome in the collection (*B. fragilis* ATCC 25285, SRA SRR4114373, against HumGut; *P. falciparum* 3D7, ENA ERR422403, against ParaRef); related is a gut metagenome (Hadza stool, ENA ERR7745403) against HumGut and *P. cynomolgi* reads (ERR035469) against ParaRef; and low is the benchmark’s cod read set against HumGut and the gut metagenome against ParaRef.

### B.4 Index construction

All SBWT indexes are first built as the bit matrix with both orientations of every *k*-mer. For the six benchmark collections, SSHash and our indexes are built from the benchmark’s eulertigs, and the one-orientation index is derived from the index with both orientations (Section 6). For HumGut and ParaRef, GGCAT [14] computes the eulertigs from the genomes (all *k*-mers kept), from which SSHash and our two-orientation index are built; the one-orientation index is built directly from the unitigs of the same *k*-mers, reoriented as in Section 6. The FBCS is converted from the one-orientation bit matrix. GGCAT’s eulertig construction is not yet exact: on ParaRef, its eulertigs lack 6 of the 54.8 · 10^9^ canonical *k*-mers of the genomes and contain 6 others twice (2 at *k* = 63). As all indexes of a collection are built from the same eulertigs, they hold the same distinct *k*-mers, and the comparisons are unaffected. Table S2 gives the construction times for HumGut and ParaRef. For each *k*-mer of a query, both SSHash and our searches report whether it is present and, if so, its identifier: the colex rank in our indexes, and SSHash’s *k*-mer ID.

**Table S2:** Construction times (wall clock) for HumGut and ParaRef. The matrices include the LCS array and the suffix-group bitvector; *^∗^*: bitvector only, no LCS array.

|  | HumGut |  | ParaRef |  |
|---|---|---|---|---|
| step | *k* = 31 | *k* = 63 | *k* = 31 | *k* = 63 |
| SSHash, from the eulertigs | 44 min | 24 min | 3 h 14 min | 51 min |
| two-or. matrix, from the eulertigs | 2 h 34 min | 6 h 20 min | 8 h 14 min | 23 h 16 min<sup>*</sup> |
| reorientation of the unitigs | 14 min | 15 min | 43 min | 31 min |
| one-or. matrix, from the unitigs | 19 min | 1 h 5 min | 3 h 45 min | 14 h 26 min<sup>*</sup> |
| one-or. FBCS, from the matrix | 35 s | 44 s | 2 min | 2 min |

### B.5 SSHash

We use the current version of SSHash (version 6, commit 85d4dfa) through a small driver written against its current API. SSHash and the driver are compiled with GCC 15.2 in CMake’s Release mode (-O3 -DNDEBUG), with SSHash’s -mbmi2 -mavx2 and with -march=native, which SSHash’s build leaves off by default. Unlike our code, SSHash is compiled without branch alignment. It is built with the benchmark’s parameters, that is, the minimizer length *m* the benchmark uses for each collection and *k*. For HumGut and ParaRef, we use *m* = 22 and 23 at *k* = 31, and *m* = 31 at *k* = 63, the largest *m* the benchmark uses (for SE and HPRC).^9^ SSHash answers reads with its streaming lookup, one query at a time, on exactly the same queries as our indexes. After each run, SSHash’s index file is dropped from the page cache, so that it never crowds our index out of the memory of the core’s NUMA node. Table S3 compares our SSHash timings with those published for the benchmark on a different CPU: they agree to within about 0.1 *µ*s on lookups and a few nanoseconds per *k*-mer on reads.

**Table S3:**
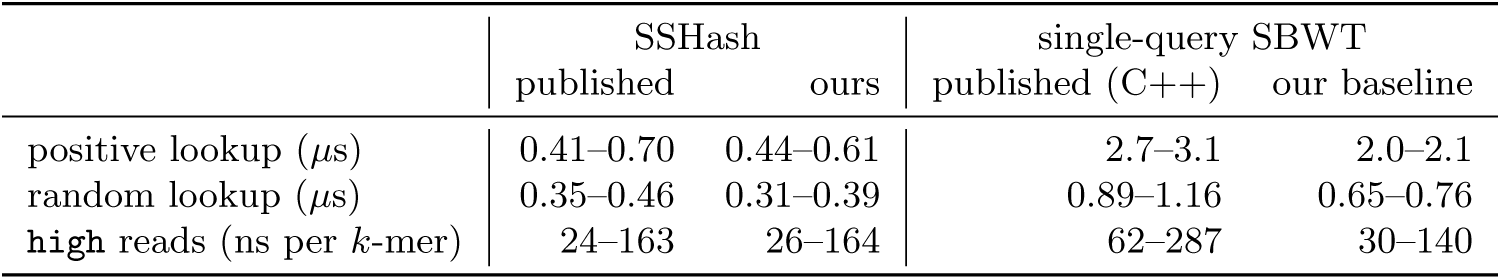
Our measurements against those published with the SSHash benchmark, *k* = 31, ranges over its six collections. The published ones were taken on an AMD Threadripper PRO 7985WX, with an earlier SSHash version, and their read timings include reading and decompressing the FASTQ files. On the left are the times for SSHash, as published by the authors and measured by us. On the right are the running times for SBWT; the times published in the SSHash benchmark paper used the SBWT-tool written in C++, and the suffix-group bitvector streaming support. Our baseline is improved with the one-line rank, and uses LCS-array streaming support.

### B.6 Code and data

The repository https://github.com/anadis504/sbwt-squeeze-rs contains our implementation of all the indexes and searches of Sections 3–7, the SSHash driver, and the scripts that build the indexes and run the experiments. All collections and read sets are public: the SSHash benchmark’s collections and read sets come from its repository, and the sources of our own collections and read sets are given in Appendices B.2 and B.3.

## C Additional results

### C.1 Performance of SBWT variants

Table S4 gives the *k*-mer lookups on the three test collections behind Section 8.2, and Table S5 the streaming queries behind Figure 5.

**Table S4:** *k*-mer lookups on the test collections, ns per *k*-mer: the single-query search and the batched search (batches of 32); 10^6^ positive *k*-mers sampled from the index and 10^6^ random *k*-mers. On the one-orientation indexes, both orientations of each *k*-mer are searched.

|  | single |  | batched |  |
|---|---|---|---|---|
| index | positive | random | positive | random |
| E. coli |  |  |  |  |
| two-orientation matrix | 1869 | 569 | 295 | 107 |
| one-orientation matrix | 2389 | 948 | 329 | 178 |
| one-orientation FBCS | 3011 | 1168 | 1083 | 605 |
| Salmonella |  |  |  |  |
| two-orientation matrix | 1957 | 616 | 299 | 117 |
| one-orientation matrix | 2502 | 1144 | 343 | 200 |
| one-orientation FBCS | 3257 | 1407 | 958 | 575 |
| human (ours) |  |  |  |  |
| two-orientation matrix | 2043 | 735 | 320 | 140 |
| one-orientation matrix | 2910 | 1355 | 375 | 242 |
| one-orientation FBCS | 4707 | 2193 | 1339 | 802 |

#### Streaming search

Every substitution shortens the matches, so the baseline slows from 95–118 ns per *k*-mer on exact reads to 144–174 ns at 5% substitutions. Batching with the LCS array gains 4.7–5.2 times on exact reads but only 2.4–2.7 times at 5%, where matches stay short and the search backs off often, reading the LCS array each time. `Forward-skip` on the bitvector is within 7% of the LCS array on exact reads. It becomes the faster of the two as substitutions increase, from 0.5% on E. coli and Salmonella, 2% on HumGut and 5% on human and ParaRef, as it decides the windows around a substitution without searching them; at 5% it is 1.2–1.6 times faster, and on random reads 3–8 times. For comparison, Table S5 also gives batched *simple k*-mer lookups with skipping. Every window is restarted and the read is searched from right to left, allowing each failed search to skip the windows to its left. Simple *k*-mer search is the fastest method on random reads, which contain no *k*-mers of the index, 1.5–1.7 times faster than `forward-skip`. On exact reads they are 13–14 times slower, as every window takes all *k* steps.

**Table S5:**
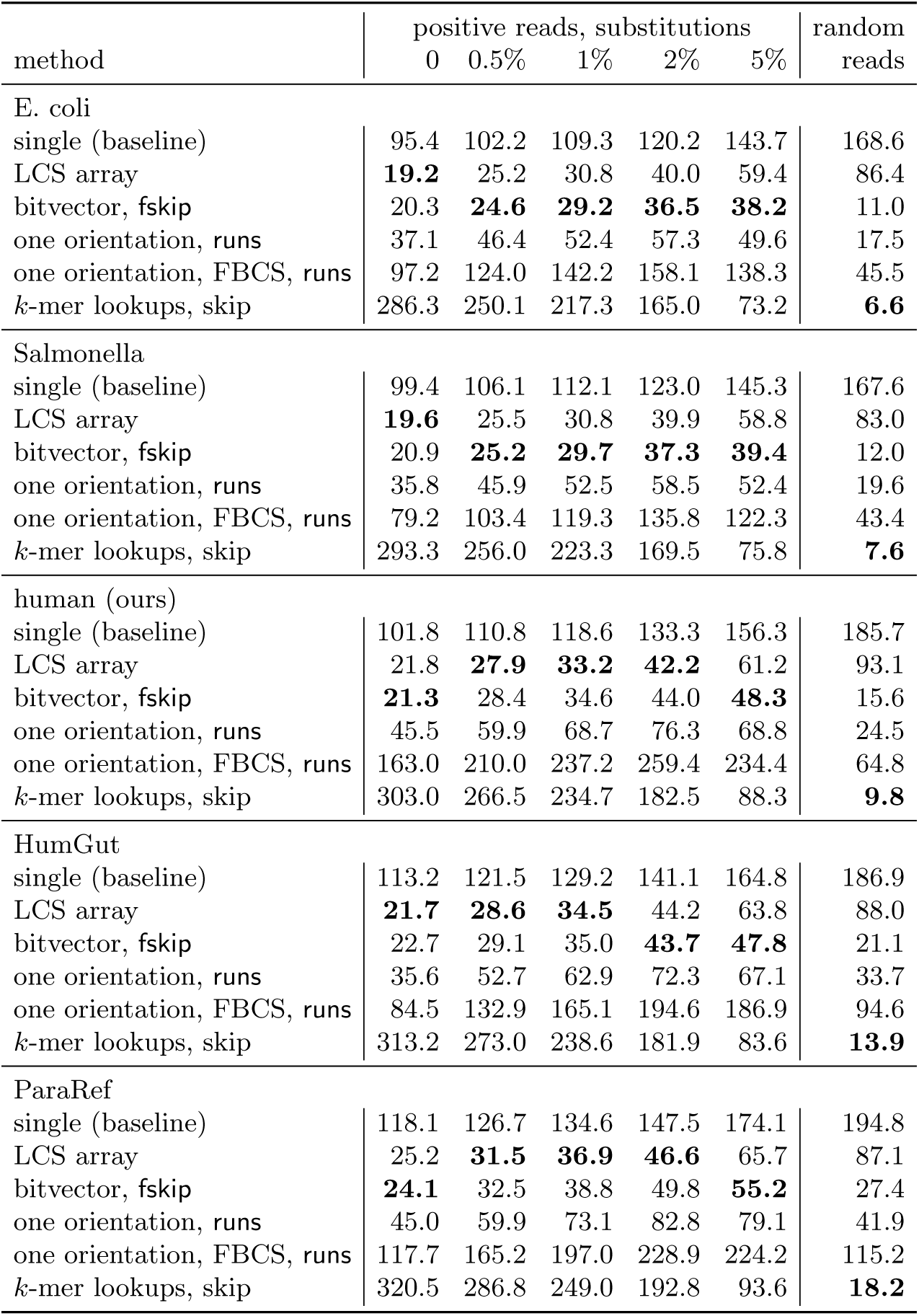
Streaming search on the test indexes, HumGut and ParaRef (*k* = 31), ns per *k*-mer of the reads (one orientation: both orientations searched): reads of 200 bases, positive ones cut from the genomes with substitutions at the given rate, and random ones; membership output, one-line rank. Fastest per column in bold.

| method | positive reads, substitutions |  |  |  |  | random |
|---|---|---|---|---|---|---|
|  | 0 | 0.5% | 1% | 2% | 5% | reads |
| E. coli |  |  |  |  |  |  |
| single (baseline) | 95.4 | 102.2 | 109.3 | 120.2 | 143.7 | 168.6 |
| LCS array | **19.2** | 25.2 | 30.8 | 40.0 | 59.4 | 86.4 |
| bitvector, fskip | 20.3 | **24.6** | **29.2** | **36.5** | **38.2** | 11.0 |
| one orientation, runs | 37.1 | 46.4 | 52.4 | 57.3 | 49.6 | 17.5 |
| one orientation, FBCS, runs | 97.2 | 124.0 | 142.2 | 158.1 | 138.3 | 45.5 |
| *k*-mer lookups, skip | 286.3 | 250.1 | 217.3 | 165.0 | 73.2 | **6.6** |
| Salmonella |  |  |  |  |  |  |
| single (baseline) | 99.4 | 106.1 | 112.1 | 123.0 | 145.3 | 167.6 |
| LCS array | **19.6** | 25.5 | 30.8 | 39.9 | 58.8 | 83.0 |
| bitvector, fskip | 20.9 | **25.2** | **29.7** | **37.3** | **39.4** | 12.0 |
| one orientation, runs | 35.8 | 45.9 | 52.5 | 58.5 | 52.4 | 19.6 |
| one orientation, FBCS, runs | 79.2 | 103.4 | 119.3 | 135.8 | 122.3 | 43.4 |
| *k*-mer lookups, skip | 293.3 | 256.0 | 223.3 | 169.5 | 75.8 | **7.6** |
| human (ours) |  |  |  |  |  |  |
| single (baseline) | 101.8 | 110.8 | 118.6 | 133.3 | 156.3 | 185.7 |
| LCS array | 21.8 | **27.9** | **33.2** | **42.2** | 61.2 | 93.1 |
| bitvector, fskip | **21.3** | 28.4 | 34.6 | 44.0 | **48.3** | 15.6 |
| one orientation, runs | 45.5 | 59.9 | 68.7 | 76.3 | 68.8 | 24.5 |
| one orientation, FBCS, runs | 163.0 | 210.0 | 237.2 | 259.4 | 234.4 | 64.8 |
| *k*-mer lookups, skip | 303.0 | 266.5 | 234.7 | 182.5 | 88.3 | **9.8** |
| HumGut |  |  |  |  |  |  |
| single (baseline) | 113.2 | 121.5 | 129.2 | 141.1 | 164.8 | 186.9 |
| LCS array | **21.7** | **28.6** | **34.5** | 44.2 | 63.8 | 88.0 |
| bitvector, fskip | 22.7 | 29.1 | 35.0 | **43.7** | **47.8** | 21.1 |
| one orientation, runs | 35.6 | 52.7 | 62.9 | 72.3 | 67.1 | 33.7 |
| one orientation, FBCS, runs | 84.5 | 132.9 | 165.1 | 194.6 | 186.9 | 94.6 |
| *k*-mer lookups, skip | 313.2 | 273.0 | 238.6 | 181.9 | 83.6 | **13.9** |
| ParaRef |  |  |  |  |  |  |
| single (baseline) | 118.1 | 126.7 | 134.6 | 147.5 | 174.1 | 194.8 |
| LCS array | 25.2 | **31.5** | **36.9** | **46.6** | 65.7 | 87.1 |
| bitvector, fskip | **24.1** | 32.5 | 38.8 | 49.8 | **55.2** | 27.4 |
| one orientation, runs | 45.0 | 59.9 | 73.1 | 82.8 | 79.1 | 41.9 |
| one orientation, FBCS, runs | 117.7 | 165.2 | 197.0 | 228.9 | 224.2 | 115.2 |
| *k*-mer lookups, skip | 320.5 | 286.8 | 249.0 | 192.8 | 93.6 | **18.2** |

#### FBCS overflow

Table S6 gives the share of FBCS blocks that overflow (Section 7). It depends on how often empty and multi-character subsets occur, and so on the collection: from under 0.1% on kestrel and NCBI-virus at *k* = 63 to 35% on SE at *k* = 31, and it is lower at *k* = 63 than at *k* = 31 on every collection.

### C.2 Comparison with SSHash

Tables S7 and S8 give all values behind Figure 6 and Table 2, and Table S9 gives lookups of mixed present and absent *k*-mers.

**Table S6:** Percentage of the blocks of the one-orientation FBCS that overflow, per collection.

| our collections |  |  | SSHash benchmark |  |  |
|---|---|---|---|---|---|
|  | *k* = 31 | *k* = 63 |  | *k* = 31 | *k* = 63 |
| E. coli | 20.2 | — | cod | 2.5 | 0.5 |
| Salmonella | 5.4 | — | kestrel | 0.4 | 0.05 |
| human (ours) | 9.9 | — | human | 5.9 | 0.9 |
| HumGut | 9.8 | 3.1 | NCBI-virus | 0.2 | 0.05 |
| ParaRef | 3.1 | 0.4 | SE | 34.8 | 8.9 |
|  |  |  | HPRC | 15.0 | 6.3 |

#### Streaming support

On high reads at *k* = 31, the LCS array and `forward-skip` are within 8% of each other. On low reads, `forward-skip` is 2.8–6.0 times faster at *k* = 31 and up to 21 times faster at *k* = 63 (HPRC), as it decides most windows without a search. Batched *k*-mer lookups with skipping, which need no streaming support, come within about 1.5 times of `forward-skip` on low reads, and are the fastest on NCBI-virus at *k* = 63, but are 6.5–27 times slower on high reads, where every window needs a restart.

#### Small and large collections

SSHash comes closest on cod and kestrel, the smallest genomes: at *k* = 31 our fastest search is only 1.1–1.3 times faster on their high reads, and at *k* = 63 SSHash is the fastest on both of kestrel’s high and related reads. On the larger and more varied collections SE and HPRC, our fastest search is 1.3–6.6 times faster on high reads.

#### *k*-mer lookups

On random *k*-mers, the two-orientation matrix is the fastest index everywhere, 2.1–2.7 times faster than SSHash at *k* = 31 and 2.4–3.2 times at *k* = 63: a random *k*-mer fails after about 17–20 characters, so its cost hardly grows with *k* (122–248 ns at both *k*). A positive *k*-mer takes all *k* steps, and its lookup time on the two-orientation matrix grows 2.3–2.5 times from *k* = 31 to *k* = 63, while SSHash’s grows 1.4–1.7 times. So at *k* = 31 the two-orientation matrix is 1.5–1.9 times faster than SSHash on positive *k*-mers, while at *k* = 63 SSHash is up to 12% faster on four of the eight collections, about as fast on human, and slower on SE, HPRC and HumGut. The one-orientation matrix, which searches both orientations of each query, is as fast as SSHash or faster on positive *k*-mers at *k* = 31 (up to 1.6 times faster), up to 1.2 times slower at *k* = 63 (but faster on SE, HPRC and HumGut), and faster than SSHash on random *k*-mers at both *k*. FBCS takes 2.0–3.0 times SSHash’s time on positive *k*-mers and 1.2–2.8 times on random ones.

#### Mixed lookups

Positive and random *k*-mers are two extremes: a random *k*-mer fails after a few characters, and its minimizer is usually absent from SSHash’s index. Table S9 gives lookups of 10^6^ *k*-mers sampled from each collection’s related reads, mixing present *k*-mers with near misses that fail after varying lengths. The two-orientation matrix is the fastest on every collection at *k* = 31 (1.2–2.4 times faster than SSHash) and on the six benchmark collections at *k* = 63 (1.3–2.3 times); SSHash is the fastest on HumGut and ParaRef at *k* = 63, by 1.1 times. On human at *k* = 31, SSHash is slower on the mix than on positive or random *k*-mers alone (538 against 515 and 370 ns): a near miss from a related genome usually has its minimizer in the index, so SSHash has to search its bucket.

**Table S7:**
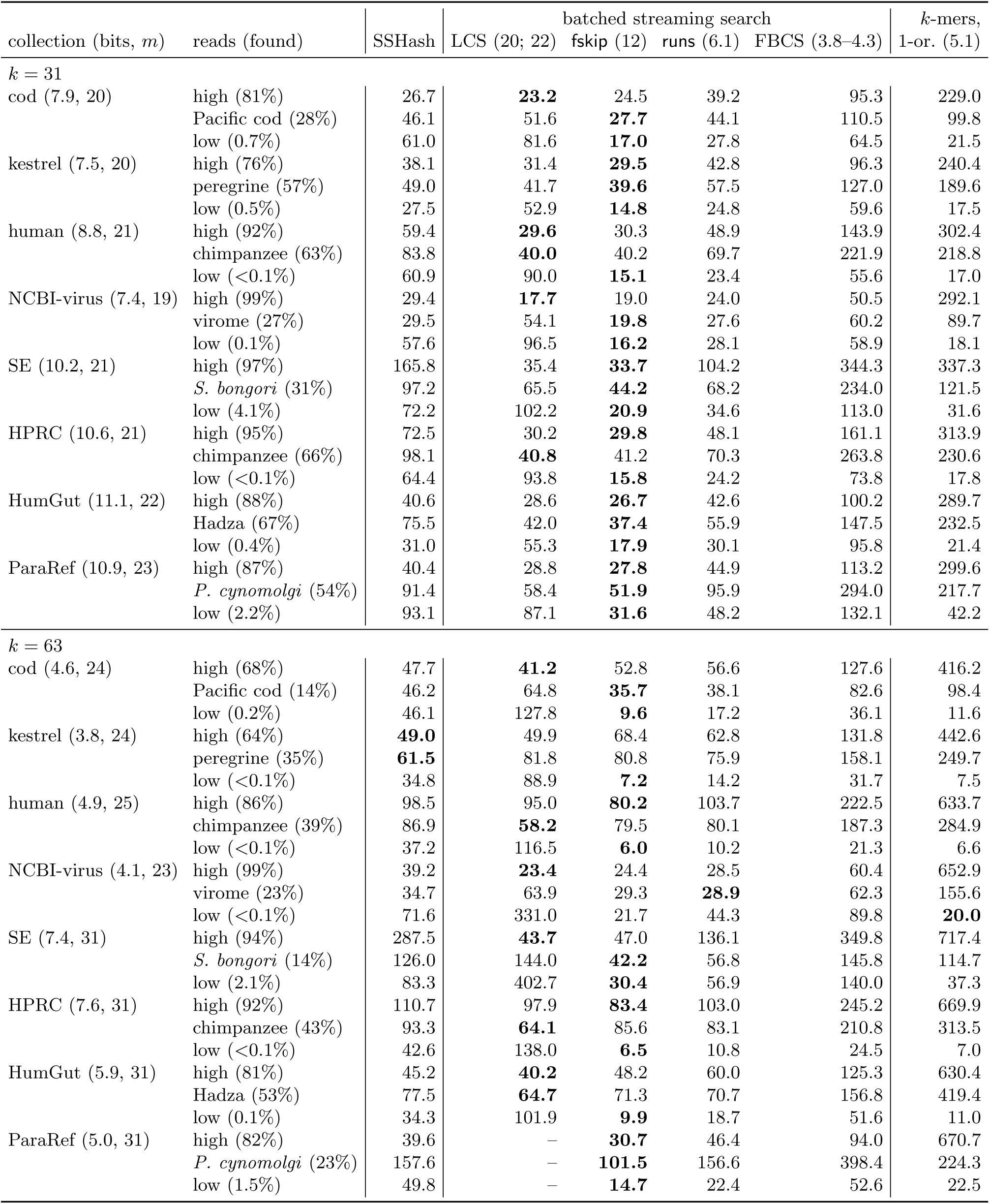
Streaming search on all collections, ns per *k*-mer of the reads, with the share of the *k*-mers found; in brackets the bits per *k*-mer of SSHash (and its *m*) and of our indexes with their streaming support (LCS array: 20 bits at *k* = 31, 22 at *k* = 63; ParaRef at *k* = 63 has none). Fastest per row in bold.

|  |  |  | batched streaming search |  |  |  | *k*-mers, |
|---|---|---|---|---|---|---|---|
| collection (bits, *m*) | reads (found) | SSHash | LCS (20; 22) | fskip (12) | runs (6.1) | FBCS (3.8–4.3) | 1-or. (5.1) |
| *k* = 31 |  |  |  |  |  |  |  |
| cod (7.9, 20) | high (81%) | 26.7 | **23.2** | 24.5 | 39.2 | 95.3 | 229.0 |
|  | Pacific cod (28%) | 46.1 | 51.6 | **27.7** | 44.1 | 110.5 | 99.8 |
|  | low (0.7%) | 61.0 | 81.6 | **17.0** | 27.8 | 64.5 | 21.5 |
| kestrel (7.5, 20) | high (76%) | 38.1 | 31.4 | **29.5** | 42.8 | 96.3 | 240.4 |
|  | peregrine (57%) | 49.0 | 41.7 | **39.6** | 57.5 | 127.0 | 189.6 |
|  | low (0.5%) | 27.5 | 52.9 | **14.8** | 24.8 | 59.6 | 17.5 |
| human (8.8, 21) | high (92%) | 59.4 | **29.6** | 30.3 | 48.9 | 143.9 | 302.4 |
|  | chimpanzee (63%) | 83.8 | **40.0** | 40.2 | 69.7 | 221.9 | 218.8 |
|  | low (<0.1%) | 60.9 | 90.0 | **15.1** | 23.4 | 55.6 | 17.0 |
| NCBI-virus (7.4, 19) | high (99%) | 29.4 | **17.7** | 19.0 | 24.0 | 50.5 | 292.1 |
|  | virome (27%) | 29.5 | 54.1 | **19.8** | 27.6 | 60.2 | 89.7 |
|  | low (0.1%) | 57.6 | 96.5 | **16.2** | 28.1 | 58.9 | 18.1 |
| SE (10.2, 21) | high (97%) | 165.8 | 35.4 | **33.7** | 104.2 | 344.3 | 337.3 |
|  | *S. bongori* (31%) | 97.2 | 65.5 | **44.2** | 68.2 | 234.0 | 121.5 |
|  | low (4.1%) | 72.2 | 102.2 | **20.9** | 34.6 | 113.0 | 31.6 |
| HPRC (10.6, 21) | high (95%) | 72.5 | 30.2 | **29.8** | 48.1 | 161.1 | 313.9 |
|  | chimpanzee (66%) | 98.1 | **40.8** | 41.2 | 70.3 | 263.8 | 230.6 |
|  | low (<0.1%) | 64.4 | 93.8 | **15.8** | 24.2 | 73.8 | 17.8 |
| HumGut (11.1, 22) | high (88%) | 40.6 | 28.6 | **26.7** | 42.6 | 100.2 | 289.7 |
|  | Hadza (67%) | 75.5 | 42.0 | **37.4** | 55.9 | 147.5 | 232.5 |
|  | low (0.4%) | 31.0 | 55.3 | **17.9** | 30.1 | 95.8 | 21.4 |
| ParaRef (10.9, 23) | high (87%) | 40.4 | 28.8 | **27.8** | 44.9 | 113.2 | 299.6 |
|  | *P. cynomolgi* (54%) | 91.4 | 58.4 | **51.9** | 95.9 | 294.0 | 217.7 |
|  | low (2.2%) | 93.1 | 87.1 | **31.6** | 48.2 | 132.1 | 42.2 |
| *k* = 63 |  |  |  |  |  |  |  |
| cod (4.6, 24) | high (68%) | 47.7 | **41.2** | 52.8 | 56.6 | 127.6 | 416.2 |
|  | Pacific cod (14%) | 46.2 | 64.8 | **35.7** | 38.1 | 82.6 | 98.4 |
|  | low (0.2%) | 46.1 | 127.8 | **9.6** | 17.2 | 36.1 | 11.6 |
| kestrel (3.8, 24) | high (64%) | **49.0** | 49.9 | 68.4 | 62.8 | 131.8 | 442.6 |
|  | peregrine (35%) | **61.5** | 81.8 | 80.8 | 75.9 | 158.1 | 249.7 |
|  | low (<0.1%) | 34.8 | 88.9 | **7.2** | 14.2 | 31.7 | 7.5 |
| human (4.9, 25) | high (86%) | 98.5 | 95.0 | **80.2** | 103.7 | 222.5 | 633.7 |
|  | chimpanzee (39%) | 86.9 | **58.2** | 79.5 | 80.1 | 187.3 | 284.9 |
|  | low (<0.1%) | 37.2 | 116.5 | **6.0** | 10.2 | 21.3 | 6.6 |
| NCBI-virus (4.1, 23) | high (99%) | 39.2 | **23.4** | 24.4 | 28.5 | 60.4 | 652.9 |
|  | virome (23%) | 34.7 | 63.9 | 29.3 | **28.9** | 62.3 | 155.6 |
|  | low (<0.1%) | 71.6 | 331.0 | 21.7 | 44.3 | 89.8 | **20.0** |
| SE (7.4, 31) | high (94%) | 287.5 | **43.7** | 47.0 | 136.1 | 349.8 | 717.4 |
|  | *S. bongori* (14%) | 126.0 | 144.0 | **42.2** | 56.8 | 145.8 | 114.7 |
|  | low (2.1%) | 83.3 | 402.7 | **30.4** | 56.9 | 140.0 | 37.3 |
| HPRC (7.6, 31) | high (92%) | 110.7 | 97.9 | **83.4** | 103.0 | 245.2 | 669.9 |
|  | chimpanzee (43%) | 93.3 | **64.1** | 85.6 | 83.1 | 210.8 | 313.5 |
|  | low (<0.1%) | 42.6 | 138.0 | **6.5** | 10.8 | 24.5 | 7.0 |
| HumGut (5.9, 31) | high (81%) | 45.2 | **40.2** | 48.2 | 60.0 | 125.3 | 630.4 |
|  | Hadza (53%) | 77.5 | **64.7** | 71.3 | 70.7 | 156.8 | 419.4 |
|  | low (0.1%) | 34.3 | 101.9 | **9.9** | 18.7 | 51.6 | 11.0 |
| ParaRef (5.0, 31) | high (82%) | 39.6 | – | **30.7** | 46.4 | 94.0 | 670.7 |
|  | *P. cynomolgi* (23%) | 157.6 | – | **101.5** | 156.6 | 398.4 | 224.3 |
|  | low (1.5%) | 49.8 | – | **14.7** | 22.4 | 52.6 | 22.5 |

**Table S8:** *k*-mer lookups on all collections, ns per *k*-mer: 10^6^ positive *k*-mers (half of them reverse-complemented) and 10^6^ random *k*-mers, the same for all indexes; bits per *k*-mer of the index alone. Fastest per row in bold.

| collection | SSHash |  |  | two-or. matrix (10) |  | one-or. matrix (5.1) |  | one-or. FBCS |  |  |
|---|---|---|---|---|---|---|---|---|---|---|
|  | bits, *m* | pos. | random | pos. | random | pos. | random | bits | pos. | random |
| *k* = 31 |  |  |  |  |  |  |  |  |  |  |
| cod | 7.9, 20 | 460 | 319 | **302** | **123** | 346 | 211 | 2.85 | 938 | 568 |
| kestrel | 7.5, 20 | 456 | 345 | **303** | **127** | 355 | 220 | 2.80 | 962 | 591 |
| human | 8.8, 21 | 515 | 370 | **312** | **135** | 366 | 235 | 2.90 | 1160 | 685 |
| NCBI-virus | 7.4, 19 | 438 | 312 | **301** | **122** | 338 | 207 | 2.80 | 856 | 543 |
| SE | 10.2, 21 | 584 | 348 | **304** | **128** | 360 | 224 | 3.27 | 1543 | 949 |
| HPRC | 10.6, 21 | 610 | 391 | **327** | **143** | 374 | 245 | 3.04 | 1496 | 901 |
| HumGut | 11.1, 22 | 611 | 417 | **360** | **178** | 419 | 314 | 2.91 | 1418 | 1125 |
| ParaRef | 10.9, 23 | 628 | 444 | **392** | **212** | 451 | 365 | 2.86 | 1512 | 1263 |
| *k* = 63 |  |  |  |  |  |  |  |  |  |  |
| cod | 4.6, 24 | **667** | 483 | 712 | **157** | 759 | 247 | 2.83 | 1892 | 606 |
| kestrel | 3.8, 24 | **630** | 509 | 708 | **160** | 756 | 255 | 2.80 | 1871 | 625 |
| human | 4.9, 25 | 732 | 541 | **730** | **169** | 785 | 270 | 2.82 | 2016 | 662 |
| NCBI-virus | 4.1, 23 | **643** | 482 | 708 | **155** | 734 | 243 | 2.80 | 1755 | 588 |
| SE | 7.4, 31 | 1002 | 514 | **717** | **162** | 814 | 263 | 3.13 | 2456 | 797 |
| HPRC | 7.6, 31 | 921 | 575 | **801** | **182** | 824 | 288 | 2.93 | 2413 | 756 |
| HumGut | 5.9, 31 | 907 | 587 | **849** | **215** | 890 | 356 | 2.86 | 2312 | 1022 |
| ParaRef | 5.0, 31 | **849** | 603 | 920 | **253** | 938 | 405 | 2.83 | 2386 | 1137 |

**Table S9:** Mixed *k*-mer lookups, ns per *k*-mer: 10^6^ *k*-mers sampled from the reads of a related organism (the related read set of each collection), present and absent ones mixed, the same for all indexes; found: the share of them in the collection. Fastest per row in bold.

| collection | *k*-mers of | found | SSHash | two-or.<br>matrix | one-or.<br>matrix | one-or.<br>FBCS |
|---|---|---|---|---|---|---|
| *k* = 31 |  |  |  |  |  |  |
| cod | Pacific cod | 28% | 338 | **183** | 266 | 828 |
| kestrel | peregrine | 57% | 430 | **244** | 326 | 910 |
| human | chimpanzee | 63% | 538 | **261** | 352 | 1293 |
| NCBI-virus | virome | 27% | 187 | **158** | 230 | 648 |
| SE | *S. bongori* | 31% | 480 | **203** | 305 | 1384 |
| HPRC | chimpanzee | 66% | 591 | **278** | 362 | 1637 |
| HumGut | Hadza | 67% | 421 | **281** | 371 | 1392 |
| ParaRef | *P. cynomolgi* | 54% | 464 | **308** | 417 | 1590 |
| *k* = 63 |  |  |  |  |  |  |
| cod | Pacific cod | 14% | 482 | **305** | 390 | 1024 |
| kestrel | peregrine | 35% | 578 | **447** | 543 | 1387 |
| human | chimpanzee | 39% | 694 | **492** | 608 | 1643 |
| NCBI-virus | virome | 23% | 362 | **286** | 365 | 926 |
| SE | *S. bongori* | 14% | 708 | **314** | 459 | 1486 |
| HPRC | chimpanzee | 43% | 769 | **557** | 655 | 1924 |
| HumGut | Hadza | 53% | **473** | 539 | 653 | 1848 |
| ParaRef | *P. cynomolgi* | 23% | **538** | 566 | 638 | 1959 |

## Footnotes

1 The last-level cache is the level closest to DRAM, typically L3.

2 SPIDER and BiRank store a smaller count relative to a superblock or to the middle of the line, at 3.3–3.8% overhead, but then need more work per query.

3 github.com/jermp/kmer sets benchmark, Zenodo record 17582116.

4 arken.nmbu.no/∼larssn/humgut

5 `-C llvm-args=-x86-branches-within-32B-boundaries`

6 github.com/human-pangenomics

7 arken.nmbu.no/∼larssn/humgut

8 github.com/jermp/kmer sets benchmark, Zenodo record 17582116.

9 On HumGut at *k* = 31, the neighbouring values of *m* did not give a clearly better index: with *m* = 23 and 24, SSHash is 6 and 14% larger and up to 6 and 10% slower on reads, and only 1.5 and 2.6% faster on positive lookups; with *m* = 21 it is 5% smaller and up to 4% faster on reads, but 2% slower on lookups.

